# RMalign: an RNA structural alignment tool based on a size independent scoring function

**DOI:** 10.1101/246793

**Authors:** Jinfang Zheng, Juan Xie, Xu Hong, Shiyong Liu

## Abstract

RNA-protein 3D complex structure prediction is still challenging. Recently, a template-based approach PRIME is proposed in our team to build RNA-protein complex 3D structure models with a higher success rate than computational docking software. However, scoring function of RNA alignment algorithm SARA in PRIME is size-dependent, which limits its ability to detect templates in some cases. Herein, we developed a novel RNA 3D structural alignment approach RMalign, which is based on a size-independent scoring function RMscore. The parameter in RMscore is then optimized in randomly selected RNA pairs and phase transition points (from dissimilar to similar) are determined in another randomly selected RNA pairs. In tRNA benchmarking, the precision of RMscore is higher than that of SARAscore (0.8771 and 0.7766, respectively) with phase transition points. In balance-FSCOR benchmarking, RMalign performed as good as ESA-RNA with a non-normalized score measuring RNA structure similarity. In balance-x-FSCOR benchmarking, RMalign achieves much better than a state-of-the-art RNA 3D structural alignment approach SARA due to a size-independent scoring function. Taking the advantage of RMalign, we update our RNA-protein modeling approach PRIME to version 2.0. The PRIME2.0 significantly improves about 10% success rate than PRIME.

**Author summary:** RNA structures are important for RNA functions. With the increasing of RNA structures in PDB, RNA 3D structure alignment approaches have been developed. However, the scoring function which is used for measuring RNA structural similarity is still length dependent. This shortcoming limits its ability to detect RNA structure templates in modeling RNA structure or RNA-protein 3D complex structure. Thus, we developed a length independent scoring function RMscore to enhance the ability to detect RNA structure homologs. The benchmarking data shows that RMscore can distinct the similar and dissimilar RNA structure effectively. RMscore should be a useful scoring function in modeling RNA structures for the biological community. Based on RMscore, we develop an RNA 3D structure alignment RMalign. In both RNA structure and function classification benchmarking, RMalign obtains as good as or even better performance than the state-of-the-art approaches. With a length independent scoring function RMscore, RMalign should be useful for the modeling RNA structures. Based on above results, we update PRIME to PRIME2.0. We provide a more accurate RNA-protein 3D complex structure modeling tool PRIME2.0 which should be useful for the biological community.

## INTRODUCTION

RNA plays important roles in many biology processes such as gene regulation, subcellular location and splicing. High-throughput global mapping of RNA duplexes with near base-pair resolution reveals that RNA interacts with RNA and RNA-binding proteins using higher order architectures in living cell[1]. Most of them, though their binding sites and binding regions[2] are determined, atomic interaction details are still missing, which is the key to understand molecular mechanisms underlying the RNA-RNA or RNA-protein recognition. With the increasing RNA and RNA-protein 3D structures deposited in PDB[3], it is important to develop better bioinformatics tools to compare RNA structures, which could provide a possible way to build atomic RNA-RNA or RNA-protein interaction models by inferring RNA structural homologs with lower sequence similarity. Some RNA structure comparing approaches have been developed under a scoring strategy with a traditional sequence alignment algorithm [4-7]. In these approaches, the RNA 3D structures are represented with structural alphabet (SA) [5-7] or dihedral angles [4]. Then DP algorithm is used to align this sequence with a substitution scoring matrix. Besides, STAR3D employs a substitution scoring function which includes the RMSD, aligned stack regions and the distance [8]. In the other state-of-the-art alignment approaches, SARA applies a statistical scoring function to measure the similarity of RNA 3D structures [9, 10]; and ESA-RNA uses the geodesic distance integrating RNA sequence with 3D structure information to measure the RNA similarity[11]. Like using a geometric concept in ESA-RNA, R3D Align and FR3D employs geometric discrepancy to measure the RNA similarity[12, 13]. Similar to SARA-Coffee [14] coupling with sequence alignments, SupeRNAlign iteratively superimposes the RNA fragment structures with R3D Align algorithm and maximizes the local fit [15]. They found that R3D is scoring best among the tools without ESA-RNA in their benchmark. Based on the state-of-the-art alignment approach SARA, a template-based approach PRIME is proposed in our team to build RNA-protein complex 3D structure models, which shows a higher success rate than computational docking software. However, the scoring function of RNA alignment algorithm SARA is size-dependent, which limits its ability to detect potential templates in some cases.

In this manuscript, we introduce an RNA structural alignment approach based on RMscore which is a size independent scoring function to measure RNA structure similarity. Firstly, we reveal the liner relationship between logarithmic length of RNA and logarithmic radius of gyration (Rg) of RNA. At the same time, aligned correlation coefficient (ACC) describing the relationship between RMSD and Rg also has a complex function relation with the RNA length. Combining these function relations, a length slightly independent scoring function RMscore is determined. Then RMscore is applied to two randomly selected independent datasets to optimize parameters and determine the transition point from similar to dissimilar. With the transition point, RMscore performs outstanding than SARAscore[5, 10] in selecting similar tRNA pairs. Then based on the RMscore, we developed an RNA structural alignment method RMalign. In RNA function classification, RMalign performs as good as ESA-RNA in balance-FSCOR. However, RNAs shared the structural similarity may have different functions. So, we benchmark RMalign in RNA structural classification. In RNA structural classification, RMalign performs much better than SARA in balance-x-FSCOR. Finally, PRIME is updated to PRIME 2.0 by replacing SARA with RMalign. PRIME 2.0 improved the success rate about 10% than PRIME when it is tested in protein-RNA docking benchmark.

## MATERIAL AND METHODS

### Datasets

We downloaded RNA structure coordinates from PDB[3] website with the RNA structure containing more than one RNA chain. (http://www.rcsb.org/pdb/home/home.do). This step obtains 2557 RNA structures. Based on these RNA structures, several datasets are constructed for variable goals. PDB-3775 is constructed to explore the relationship between the Rg of RNA and the RNA length. Fragment dataset is built to study the relationship between the ACC and RMSD in RNA. Results in PDB-3775 and fragment-pairs are combined together to estimate the expression of RMscore. Because calculation of all-to-all alignments of 3775 RNAs is time consuming process, we randomly select two RNA-RNA pair datasets without overlap. Random pairs-0.3M and random pairs-0.1M are built to optimize the compensation and determine the transition point, respectively. As benchmarking modeRNA in tRNA[16], the tRNA-pairs are also created for benchmarking RMscore. Balance-FSCOR and Balance-x-FSCOR are established to benchmark RMalign in function and structure classification. At last, the unbound protein-RNA docking set is employed to compare the performance of PRIME 2.0 and PRIME.

#### PDB-3775

Total 2557 RNA structures and their complexes from PDB are separated by chains. The RNA structures in mmcif format are not kept. Finally, 3775 RNA chains are kept. PDB-3775 represents all RNA structures in PDB. The relationships between the Rg of RNA and the RNA length are explored in this dataset.

#### Fragment-pairs

ACC in proteins describing the relationship between RMSD and Rg is reported in the ref.[17]. In order to study the relationship between the ACC and RMSD in RNA, we generate a fragment pair dataset based on PDB-3775. For 3775 RNA chains, only one fragment is randomly chosen for each RNA chain. Then all the fragments with the identical length are made into pairs. This strategy generating structure fragments is previous used in the protein field[18]..

#### Random pairs-0.3M

We randomly selected 0.3 million RNA pairs from all-to-all alignment of RNA chains in PDB-3775 to optimize parameters in RMscore. The alignment of the paired RNA is generated by needle[19]. This data is named as random pairs-0.3M.

#### Random pairs-0.1M

We randomly chose 0.1 million RNA pairs from all-to-all pair of RNA chains in PDB-3775 to determine the phase transition from all-to-all pairs. The alignment of the paired RNA is aligned by SARA[5, 10], which is an RNA structural alignment protocol based on unit-vector root-mean-square. This data is named as random pairs-0.1M.

#### tRNA paris

We downloaded all tRNA structures from NDB[20] (http://ndbserver.rutgers.edu/). We extracted one RNA chain from one structure or its complex. This process outputs 175 RNA chains. tRNA pairs were then constructed through all-to-all pairwise aligned by SARA.

#### Balance-FSCOR

FSCOR[5, 10] is downloaded from this website (http://structure.biofold.org/sara/datasets.html), which is constructed from SCOR[21] to benchmark RNA structural alignments. Positive pairs are generated from the RNAs with the same function in FSCOR. Negative pairs are generated from randomly selected the RNAs with different functions. The number of negative pairs is equal to the number of positive pairs. This data including both negative pairs and positive pairs is named as balance-FSCOR.

#### balance-x-FSCOR

In protein structural alignment field structural similarity is often used as an evaluation metric. However, in RNA structural alignment field, RNA function is used as the metric in benchmarking RNA structural alignment tool on the balance-FSCOR. Through analysing the pairwise RNA structural alignment results, we found that two RNAs may have different functions even their structures are very similar. It is debatable to benchmark RNA structural alignment algorithm with RNA functions only. So, we constructed balance-x-FSCOR to benchmark RNA structural alignment approach employing the RNA structural similarity RMScore as a metric. Firstly, FSCOR is clustered by RMalign with different RMscore cut-offs (x=0.4, 0.45, 0.5 … 1.0) to construct the x-FSCOR. So, each RNA structure in x-FSCOR is assigned to a cluster. The positive pair is defined as the structures are in the same cluster and the negative pair is defined as in the different cluster. For saving time (SARA runs slowly in large structures alignment), only 1000 positive and negative pairs are randomly selected from all-to-all pairs of x-FSCOR. These datasets are named as the balance-x-FSCOR. If the number of positive or negative pairs is less than 1000, the dataset contains less pairs. The structural classes with various cut-offs of 419 RNA chains in FSCOR can be downloaded from www.rnabinding.com/RMalign/RMalign.html. The vary cut-offs are tried, because it is still unknown which value is appropriate to cluster the RNA structures.

#### Unbound protein-RNA docking set

The unbound set is used to compare the performance of PRIME[22] and PRIME(2.0) in predicting protein-RNA complex structures. This set includes 49 protein-RNA structures from protein-RNA docking benchmark[23].

### Relationship between the Rg of RNA and the RNA length

The Rg of protein is an important metric to describe the compactness of a protein. Previous studies[24, 25] based on PDB reveal a scaling law about protein Rg 
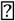
 N^0.4^ where N is the number of residues in a protein. Following in the same direction, we investigate the relationship between Rg of the RNA and its length. Simply, all RNA structures are represented with C3’ atoms. The average log Rg located in the same length bins is calculated. After calculating all RNAs in PDB-3775, we observed that a scaling law Rg 
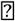
 N^0.39^ for RNA. Previous study[26] on Rg of RNA shows that this scaling law is Rg 
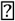
 N^1/3^. In their work, the data is fitted by a fixed index of the exponent (1/3), although Flory theory[27] limits the scaling law value between 1/3 and 3/5. Different to Hyeon *et al.*’s result[26], our study shows that the index of exponent is 0.39.

### Relationship between the ACC and the RNA length

The ACC has a function correlation with the Rg of protein and RMSD[17]. The Rg of the protein and RMSD also depend on the number of residues and the aligned length. Like protein, the relationship between RMSD of two aligned RNA structures and the Rg of these two can be written as (1) by Zhang and Skolnick[28]:

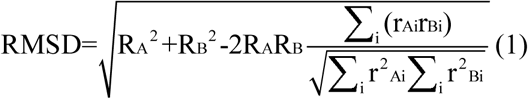

where R_A_(R_B_) is the Rg for structure A(B), rAi(rBi) is the coordinate vector after superposition.

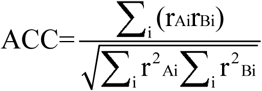

In order to reveal the relation between the ACC and aligned nucleic acids, we took a similar strategy used in TMalign[29]. Firstly, a fragment-pairs dataset is constructed. Secondly, the RMSD of RNA fragment pairs and the Rg of each fragment are calculated with the C3’ atom. Thirdly, the average and standard error of ACC are calculated if fragment pairs have the identity length. Eq. (2) reveals the function relation between the fragment length and the ACC after data fitting.

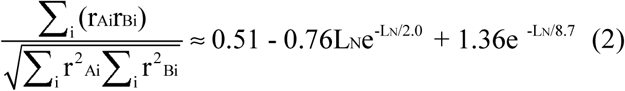

Where L_N_ is the length of fragment in the fragment-pairs.

### RMscore

Inspired by a protein 3D structure scoring function TM-score, we introduce a size-independent scoring function RMscore to describe the similarity of two RNA structures. For an RNA alignment, RMscore is defined as:

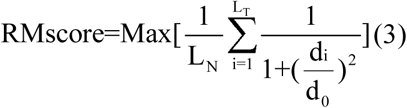

Where L_N_ is the length of average of target and query RNA, L_T_ is the length of aligned nucleic acids to the target structure, d_i_ is the distance between the ith pair of aligned nucleic acids and d_0_ is a scale to normalize the length effect. ‘Max’ denotes the maximum after optimal superposition. For different scoring strategy, a different d_0_ is adopted. In RMscore, a length-dependent d_0_ is adopted. The relation between d_0_ and the length can be estimated from Rg ∝ N^0.39^ and eq. (2).

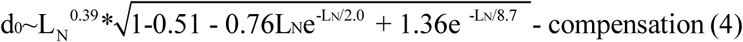

Here the constant compensation (set as 0.6) is introduced to smooth the curve when the RMscore is optimized in random pairs-0.3M (Figure S1). Eq. 4 can be well approximated by a simple formula.

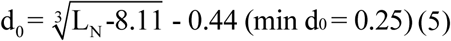

### Searching Engine of RMscore

To find the spatially optimal superposition of the query and the target structure with the maximum RMscore according to eq. (3) and eq. (5), we use an iterative searching algorithm from TM-score.

### Benchmarking RMscore on tRNA pairs

The all tRNAs are selected to benchmark RMscore because the RNA homology modelling method modeRNA was also benchmarked in tRNA dataset[16]. For comparison of the ability to select the similar RNA structures for RMscore and SARAscore (normalized SARAscore), we determine the phase transition point of RMscore and SARAscore in random pairs-0.1M. All alignments of RNA pairs are generated by SARA. The target SARAscore is normalized by dividing SARAscore of aligning itself. After the phase transition point from dissimilar to similar pairs is determined, RMscore and SARAscore are tested in tRNA pairs to distinguish similar (RMSD <= 5Å) or dissimilar (RMSD > 5Å) tRNA pairs. The alignments of tRNA pairs are also generated by SARA. A possible application of RMscore is to measure similarity between the native RNA structures and RNA models[16].

### RMalign

We developed RMalign, an RNA structural alignment tool based on RMscore. The strategy taken by RMalign is similar to TM-align (see Figure S4). Two processes are modified. Firstly, in the secondary type of initial alignment, RNA secondary structure is calculated by X3DNA[30]. Secondly, for the final scoring process, all the aligned nucleic acids are used to score instead of setting a distance cut-off for the aligned nucleic acid.

### Benchmarking RMalign on balance-FSCOR and balance-x-FSCOR

RMalign and ESA-RNA are tested on the balance-FSCOR for RNA function classification. They are also tested on the balance-x-FSCOR for RNA structural similarity.

The purpose of RNA structure alignment approach is used to detect the structural similarity. So, we also benchmark RMalign in balance-x-FSCOR. The AUC value is used as the metric to measure the performance.

### Predicting protein-RNA 3D structure

We previous developed an approach PRIME[22] to predict the protein-RNA 3D structure. PRIME was tested on an unbound protein-RNA docking benchmark. The result shows that PRIME performs better than 3dRPC[31]. We update previous PRIME to v2.0, because RMalign performs better than SARA in balance-x-FSCOR. A similar approach with PRIME is adopted to build the protein-RNA complex structure model. The transformation matrices of TM-score and RMscore are applied to superimpose the target protein and RNA onto the templates. The ligand RMSD of RNA C3’ atom between the model and the native structure is calculated. The quality of the model is measured by ligand RMSD. A prediction defined as “acceptable” for the ligand RMSD <= 10 Å[32].

## RESULTS

### Principle and benchmarking of RMscore

In figure 3(A), it shows that the raw-RMscore changes with the length. To overcome the shortcoming of length-dependent scoring function (Fig. 3A) for aligning RNA structures, we proposed a size-independent scoring function RMscore (all RMscore discussed in this manuscript is normalized by an average length) to measure the RNA structural similarity. In order to obtain the formula of RMscore like TM-score, firstly we reveal that Rg of RNA has liner relation (R^2^ = 0.91) with the logarithmic RNA length (Fig. 1). Secondly, we found that aligned correlation coefficient has a complex function relation (R^2^ = 0.95) with the number of aligned nucleic acids (Fig. 2). Then a compensation value 0.6 is introduced to flat the average RMscore in random pairs-0.3M. (Fig. S1). The final average RMscore shows a slightly dependent on the RNA length with the highest standard error 0.2 (Fig. 3B). For comparing RMscore with normalized SARAscore, the relationship between the RMSD and the RMscore/normalized SARAscore in 0.1 million RNA pairs randomly selected from total pairs are investigated. In Fig. S2, it shows that the phase transition (from noise to similar RNA pairs) are 0.5 and 0.78 (at accumulative fraction = 0.5) for RMscore and normalized SARAscore, respectively. For the ability of selecting similar RNA pairs with phase transition as the cut-off, RMscore and SARAscore discriminate 0.8771 (Fig. 4A) and 0.7766 (Fig. 4B) pairs in all-to-all pairwise structure comparison for 172 tRNA structures, respectively. The result shows the RMscore can distinguish similar (RMSD <= 5Å) or dissimilar (RMSD > 5Å) RNA pairs (Fig. S3). The above results indicate RMscore is a better metric to measure RNA structural similarity than SARAscore.

**Figure 1.**
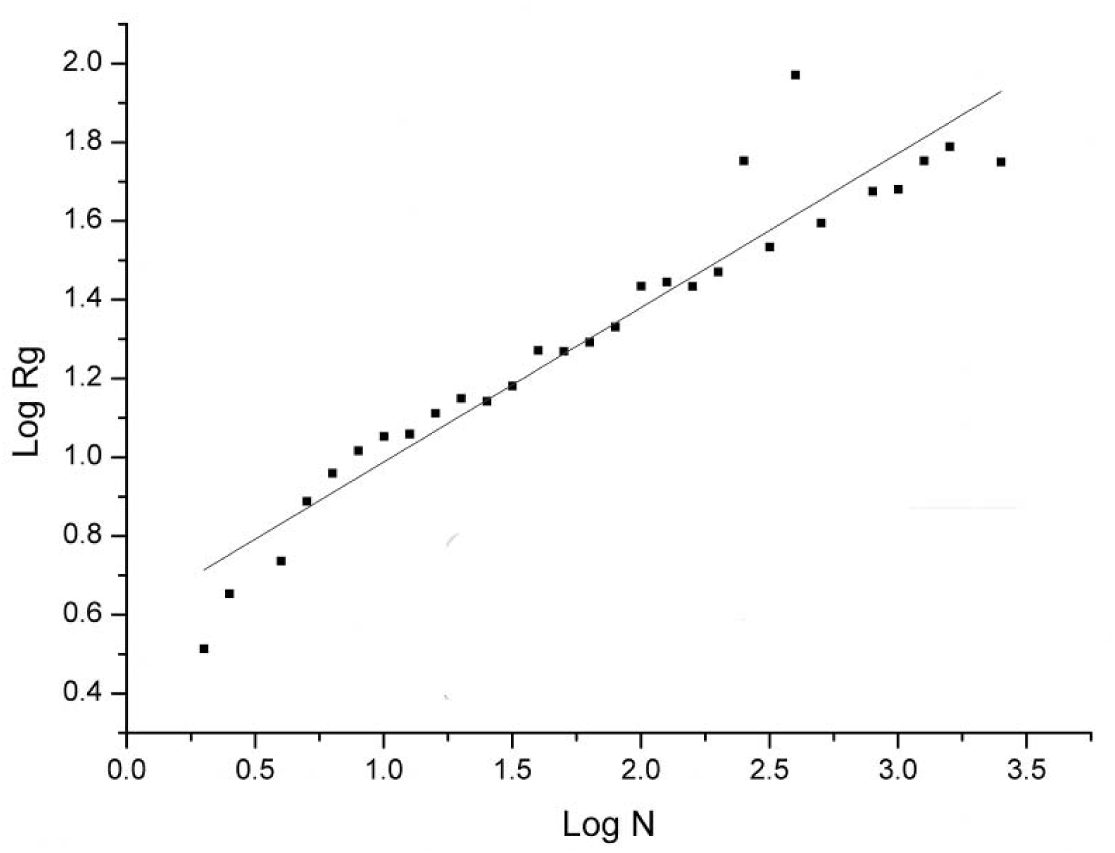
Log N vs Log Rg. The log of average Rg of RNA is plotted against log length of the corresponding RNA for 3775 RNAs. Rg is calculated with C3’ atom. The log length is split into 29 bins across the min to the max log RNA length. The Rg located in the same length bin is represented with the average value. The standard errors are not shown here. The data is fitted with a liner function (y(x) = a + b*x). Parameter of a = 0.60 ± 0.04 and b =0.39 ± 0.02.

**Figure 2.**
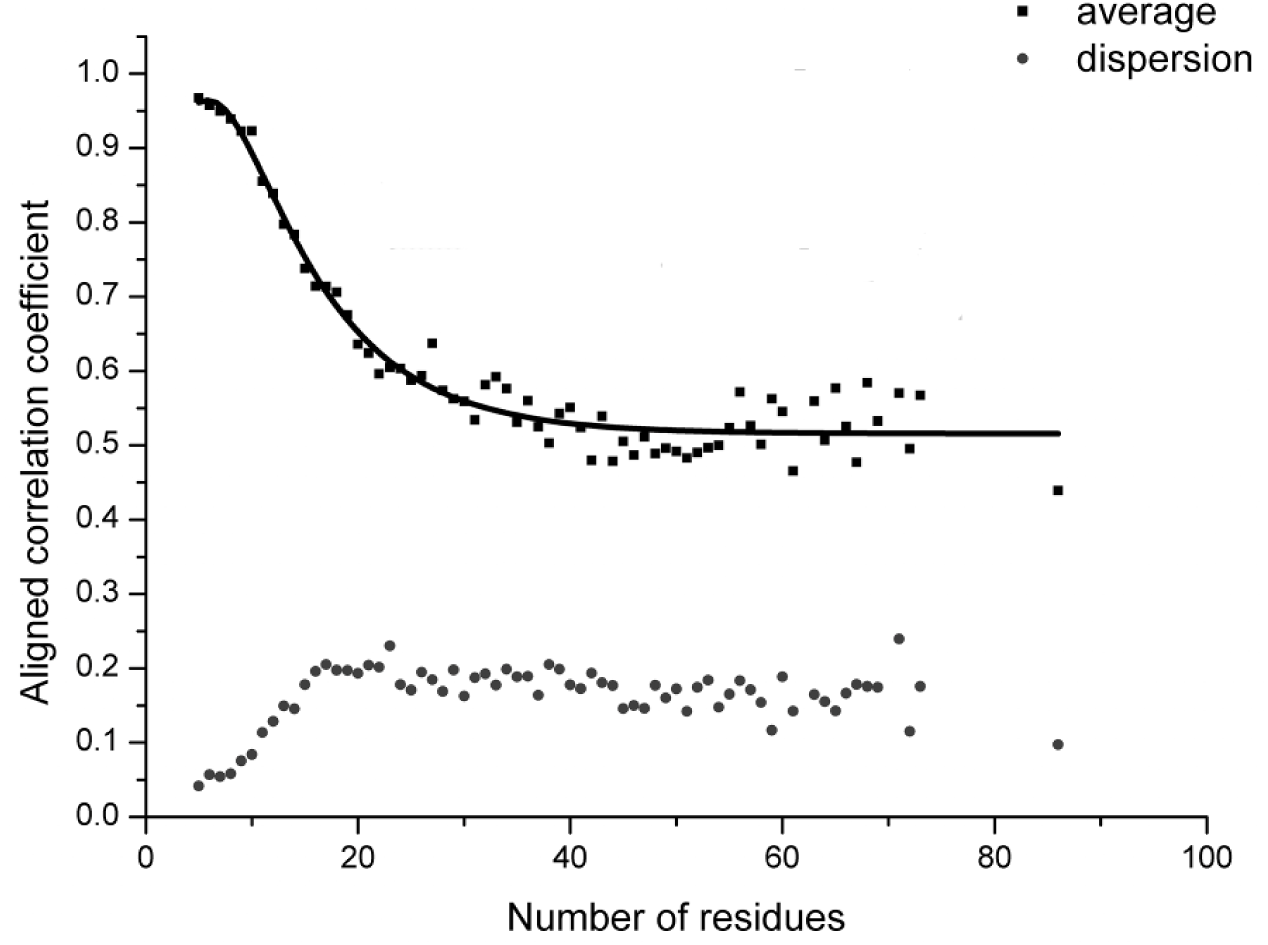
Number of residues vs aligned correlation coefficient. Average aligned correlation coefficient is plotted against number of residues. The data is calculated from RNA fragment pairs. Because the length of RNA is dispersive, the length with the number of pairs <= 20 are filtered out. The square points are the average of aligned correlation coefficient. These points are fitted with nonlinear function (y(x) = E – A*x*exp(x/B) + C*exp(x/D)). Parameter of A = 0.76 ± 0.29, B = −2.02 ± 0.57, C = 1.36 ± 0.41, D = −8.73 ± 1.30 and E = 0.52±0.01. The circle points are the standard error of aligned correlation coefficient.

**Figure 3.**
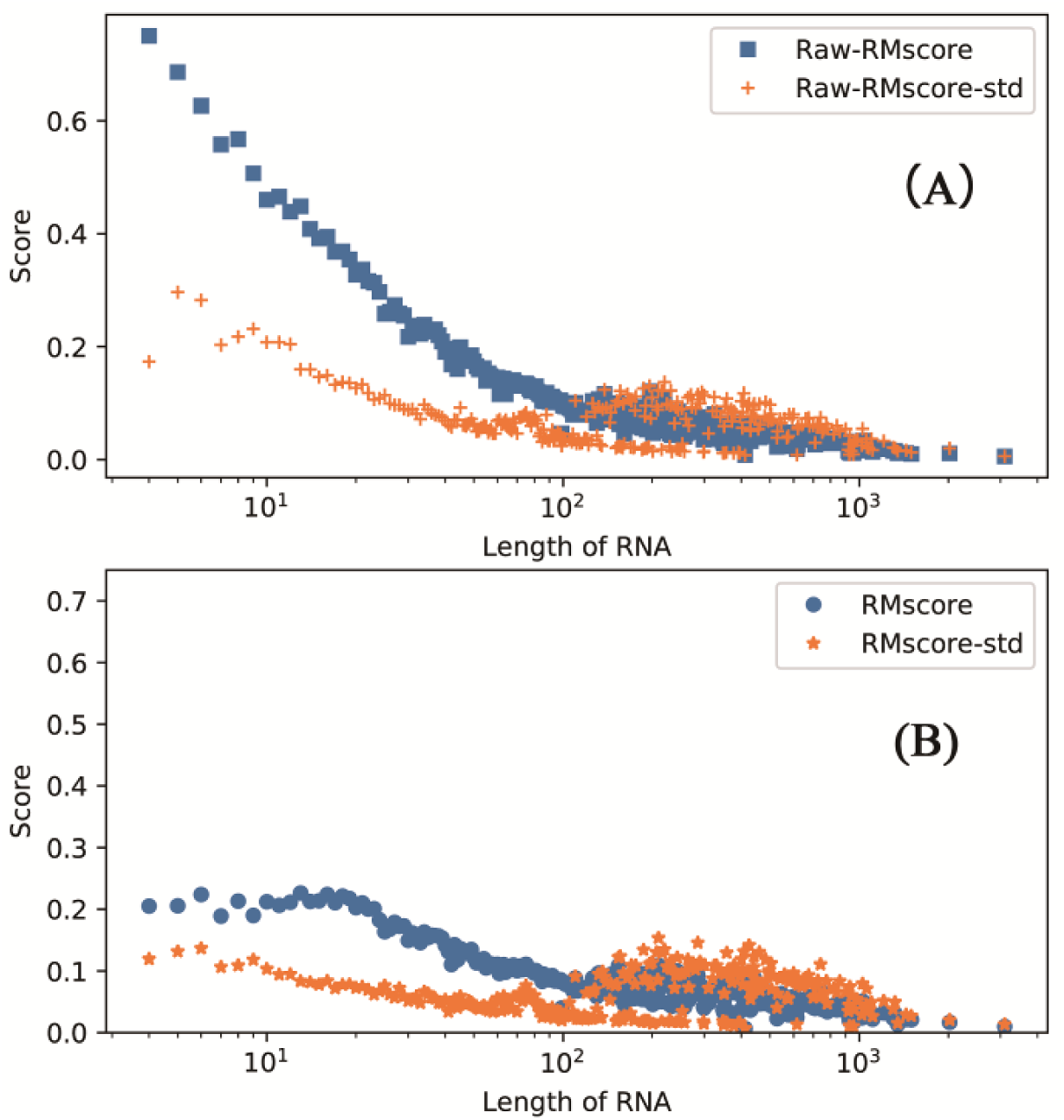
Length of RNA vs RMscore(B)/Raw-RMscore(A). The data is calculated from 0.3 million random selected pairs. RNA sequence alignment is accomplished by needle in EMBOSS package. The definition of RMscore is derived from TM-score. Raw-RMscore is calculated with the definition of RMscore but d0 = 5. The legend with a suffix “−std” are the standard error of corresponding score.

**Figure 4.**
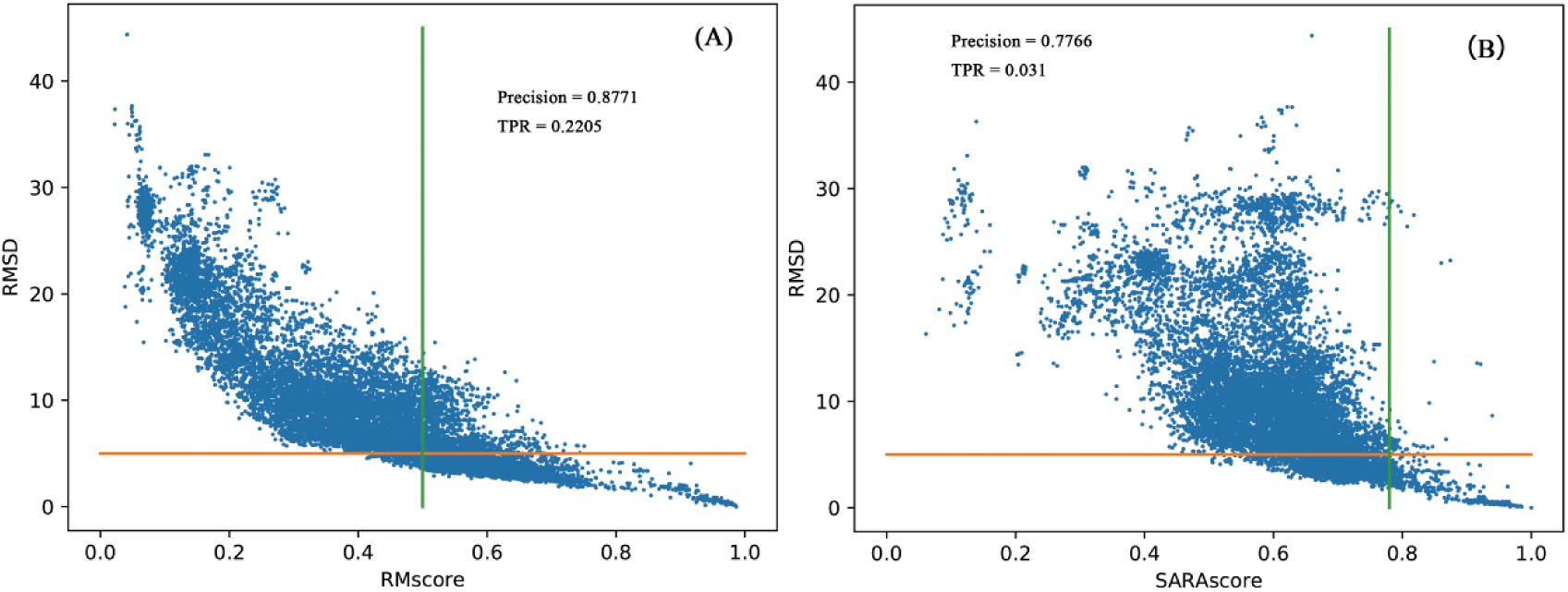
Performance of RMscore and SARAscore in identifying similar tRNA pairs. RMSD are plotted against RMscore (A) and SARAscore (B) for all-to-all pairwise comparison of 175 tRNA structures. Horizontal line (RMSD = 5Å) and vertical line (score = phase transition value at 0.5 (A) and 0.78 (B)) divided the figures into 4 quadrants. Precision is defined as that the number of points in upper left plus the number of points in bottom right divided by total number of points. The precision of RMscore (A) is 0.8771 and SARAscore (B) is 0.7766. TPR is defined as that the number of points in bottom right divided by total number of points. The TPR of RMscore is 0.2205(A) and SARAscore is 0.031(B).

### Benchmark of RMalign and comparison with other state-of-the-art approaches

For benchmarking RMalign in RNA function classification, FSCOR are downloaded and then a new dataset balance-FSCOR is constructed. RMalign obtains the AUC value of 0.95 which is as good as ESA-RNA in balance-FSCOR (Fig. 5). However, RMalign has two advantages comparing with ESA-RNA. Firstly, RMalign is written with C++ and ESA-RNA is written with a commercial software Matlab. Secondly, the geodesic distance describing the RNA structural similarity in ESA-RNA is not normalized and RMscore is a size independent score. In Fig. S5, it shows that the distribution of RMscore of negative and positive pairs in balance-FSCOR. This figure indicates that RMalign can distinguish negative and positive pairs clearly. In Fig. S6, it also shows ACC (highest value 0.88), MCC (highest value 0.73), F-measure (highest value 0.87) values of RMscore with different cut-offs. A false positive alignment example (total 13 such cases with cut-off = 0.6) shows RMalign detects the RNA structural similarity but they have the different function (Fig. S7). These 13 cases indicate that RNA function is not an appropriate metric to evaluate RNA structural alignment approach. For comparing the performance of RNA alignment tool with RNA structural similarity instead of RNA functions, we perform all-to-all alignment of FSCOR to re-cluster the 419 RNA chains with different RMscore x as cut-off. Then balance-x-FSCOR is constructed for comparing of RMalign and SARA in structural classification. In Fig. S8, it shows that the AUC of RMalign is higher than the AUC of SARA in balance-x-FSCOR. The performances on balance-FSCOR and balance-x-FSCOR show that RMalign can be used to predict RNA functions based on RNA structural similarity.

**Figure 5.**
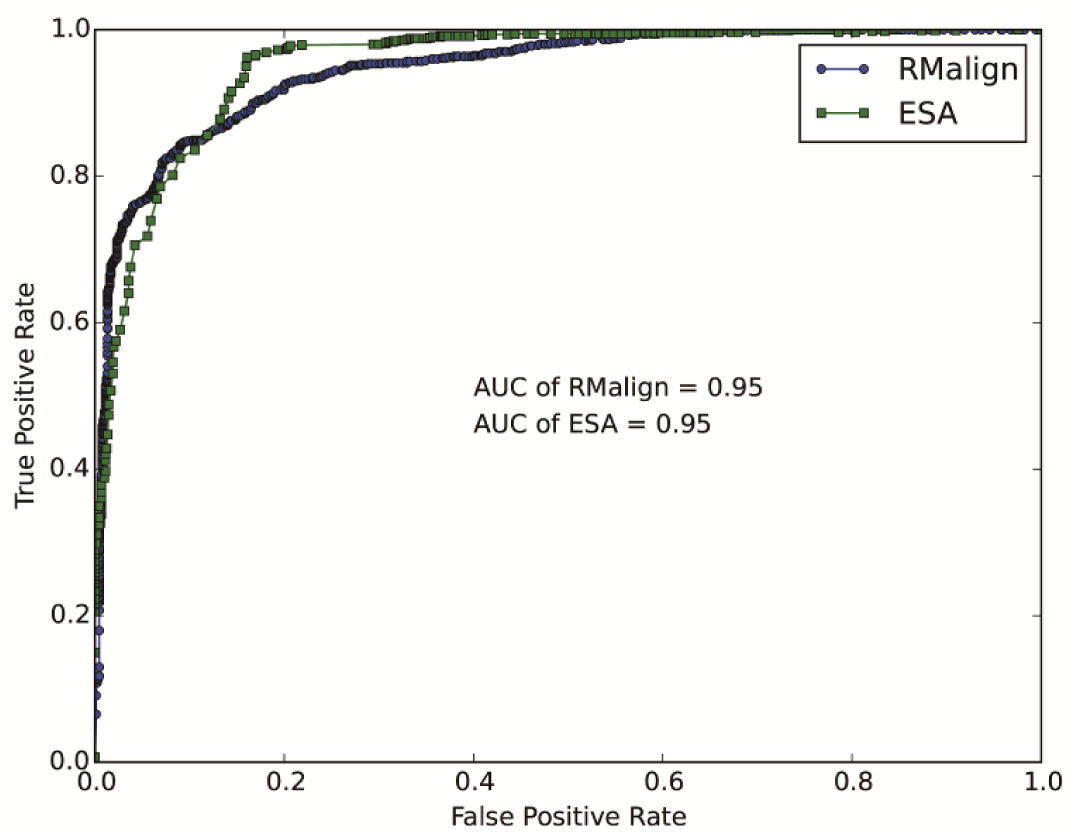
ROC curves for benchmarking in balance-FSCOR. The RNA pairs in the same functional class are regarded as the positive and pairs in the different functional class are regarded as the negative. The AUC of RMalign is 0.95 which is equal to that of ESA-RNA.

### Predicting protein-RNA 3D structure with PRIME 2.0 and comparison with PRIME

RMalign could find more remote RNA structural homologs than SARA. It could enhance searching RNA templates in the template-based RNA-protein structure modeling approach PRIME. For testing the ability to detect more potential RNA-protein complex structure templates, we update PRIME to PRIME 2.0 by replacing SARA with RMalign. PRIME 2.0 was tested on unbound RNA-protein docking benchmark containing 49 complexes. In Fig 6, it shows the RNA-protein docking results. For top 1 prediction, the success rate of PRIME 2.0 is about 10% higher than that of PRIME. The result indicates that RMscore can select more potential templates than SARAScore in protein-RNA 3D complex structure prediction. For the top 300 predictions, success rate of PRIME 2.0 is higher than PRIME. In Fig 7, it shows a successful example in PRIME 2.0 but it failed in PRIME. Above results indicate that RMalign can detect more templates for protein-RNA complex structure modeling.

**Figure 6.**
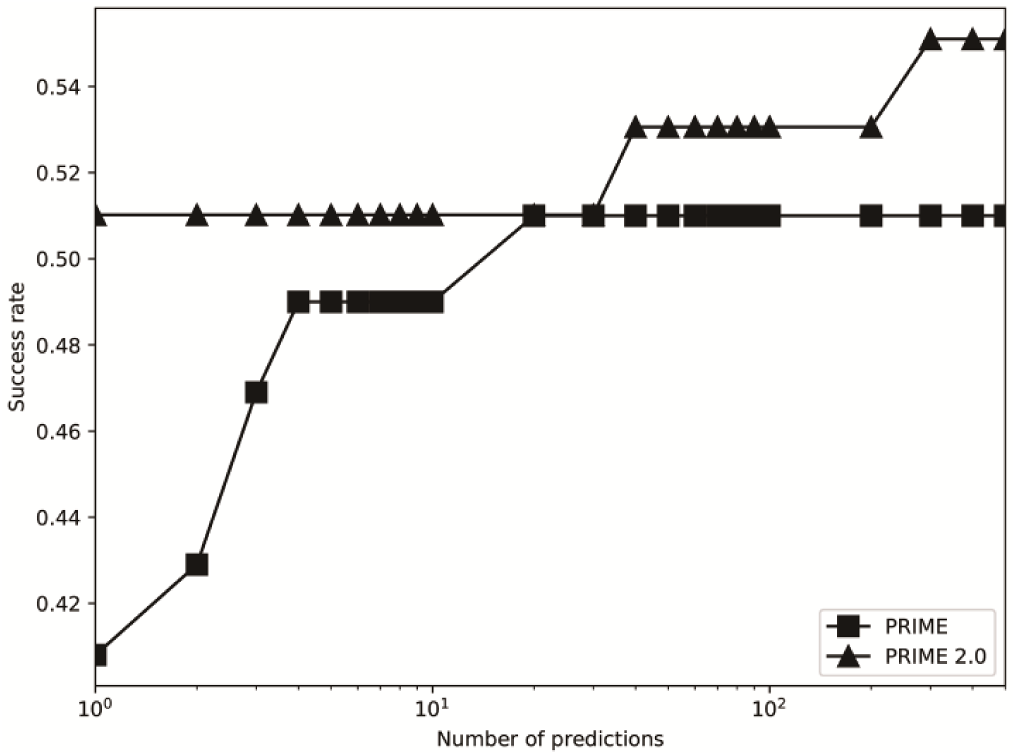
Comparison of previous PRIME. The template docking was performed by PRIME and PRIME 2.0. The successful prediction was defined as at least one match with ligand RMSD <= 10Å in top 10 and top 500 (for template docking the max prediction number is 439).

**Figure 7.**
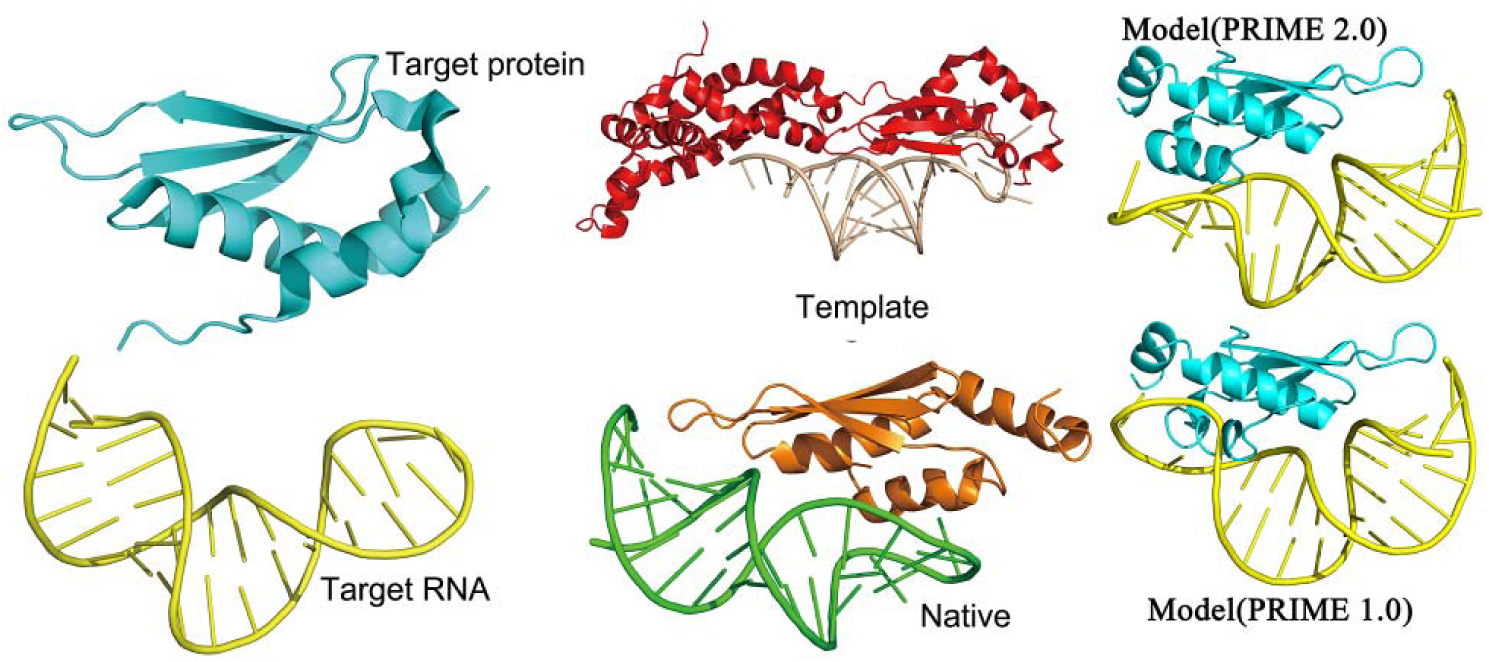
An example of target modeled by updated PRIME but PRIME fails in this case. The target, 1t4o chain A and 1t4l chain A, was modeled on the template 4oog, chain C and D. The target/template structural similarity for protein is 0.462 (TM-score) and RNA is 0.600 (RMscore). The ligand RMSD for the model (PRIME 2.0) is 6.81Å and for the model (PRIME 1.0) is 12.57 Å. The case shows that RMalign can detect more templates than SARA.

## DISCUSSION

In conclusion, we introduce an RNA structure alignment approach RMalign, which includes RMscore as the similarity score. The definition of RMscore is derived from TM-score which has been applied to protein structural alignment successfully. However, the RMscore shows a slightly dependent on RNA length. This phenomenon may be caused by the flexible structure of RNA. It is hard to benchmark RMscore like TM-score because that study in RNA falls behind in protein. For example, the best way to benchmark RMscore is to compare the similarity between RNA model and native structure in RNA structure modelling. However, no related studies have been investigated. Even more, the RNA homology modelling modeRNA employs RMSD or LG-score, which is introduced as an auxiliary metric to measure the RNA structure similarity without any modification[16]. Considering the currently situation, we study the relationship between RMscore and RMSD in RNA. The result shows that RMscore = 0.5 can discriminate the similar and dissimilar structures.

## AVAILABILITY

RMalign can be downloaded from www.rnabinding.com/RMalign/RMalign.html. PRIME 2.0 can be accessed from www.rnabinding.com/PRIME/PRIME2.0.html. All datasets used in this manuscript can be downloaded from www.rnabinding.com/RMalign/RMalign.html.

## SUPPLEMENTARY DATA

### ACKNOWLEDGEMENT

We thank the National Supercomputer Center in Guangzhou for support of computing resources.

### FUNDING

This work has been supported by the Special Program for Applied Research on Super Computation of the NSFC-Guangdong Joint Fund (the second phase) under Grant No. U1501501 and the Fundamental Research Funds for the Cen-tral Universities [2016YXMS017].

## FIGURES

**Figure S1.**
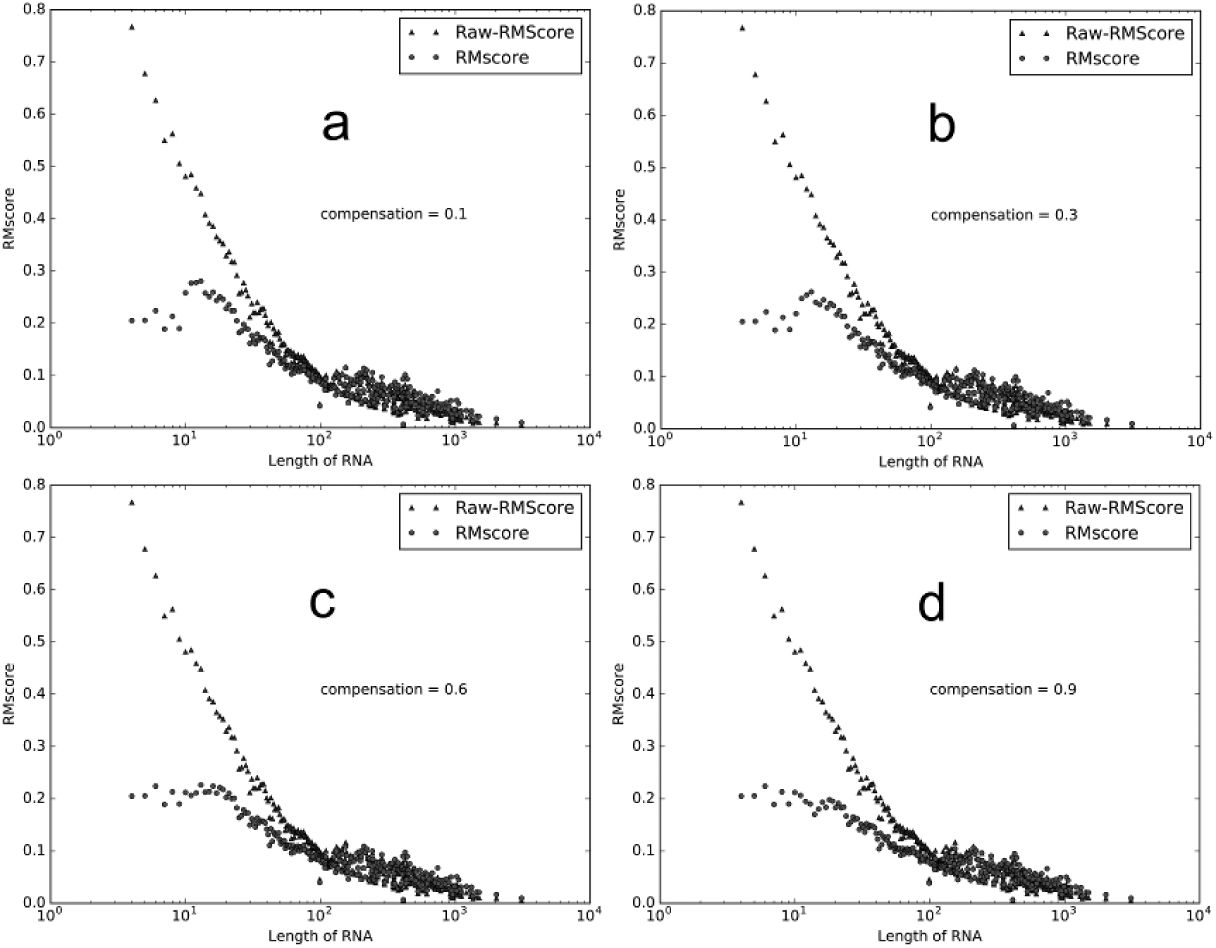
Length of RNA vs RNA structural similarity. Length of RNA is plotted against RMscore with a different compensation (a is 0.1, b is 0.3, d is 0.6 and d is 0.9) to flat the average RMscore at a smaller length, in random pairs 0.3M. RNA alignment is accomplished by needle in EMBOSS package. Raw-RMscore means that d0 is assign to 5 angstroms in RMscore definition. The compensation is chosen as 0.6 because it results in the most smoothed curve.

**Figure S2.**
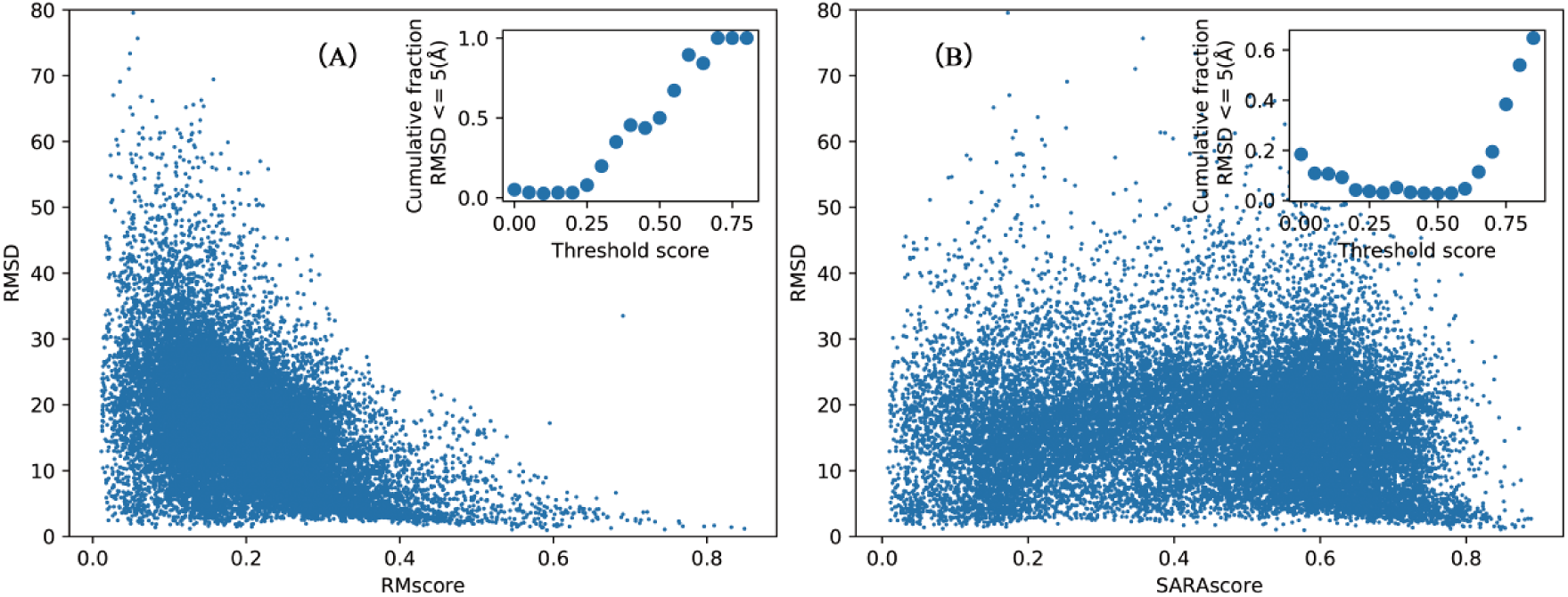
RNA structural similarity vs RMSD. SARAscore (A) and RMscore are plotted against RMSD in 0.1 million randomly selected pairs, RNA structural alignment are accomplished by SARA. The insets show the fraction of RNA-RNA pairs with RMSD <= 5 Å plotted with 0.05 bins to show the phase transition from dissimilar RNA pairs to the similar pairs.

**Figure S3.**
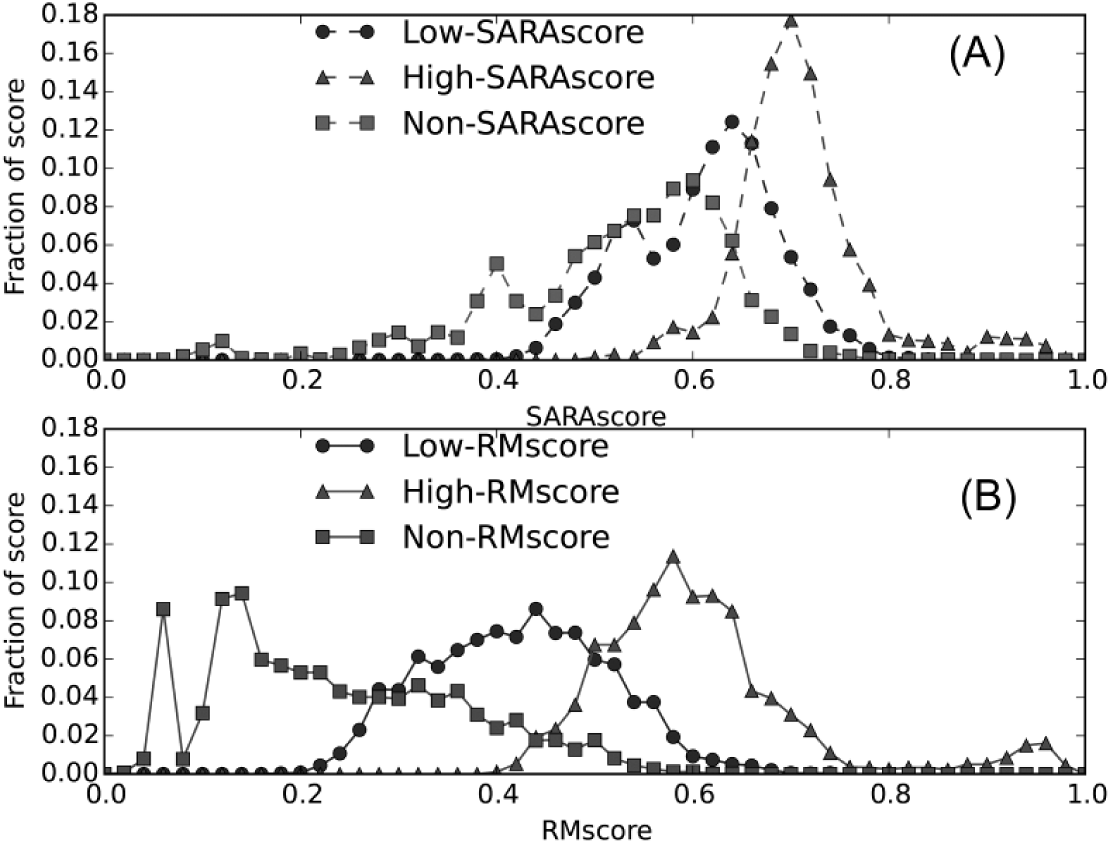
Distribution of RMScore (B) and SARAscore (A). All-to-all pairs are categorized into Low-RMScore (5Å < RMSD <= 10Å), High-RMscore (RMSD <= 5Å) or Non-RMscore (RMSD >10Å). The same category criterion is applied for SARAscore. The peak generating by RMscore are separated clear, which can be used to distinguish similarity or dissimilarity RNA pairs. However, the peak generating by SARAscore are pretty close, which is hardly used to distinguish similarity or dissimilarity RNA pairs.

**Figure S4.**
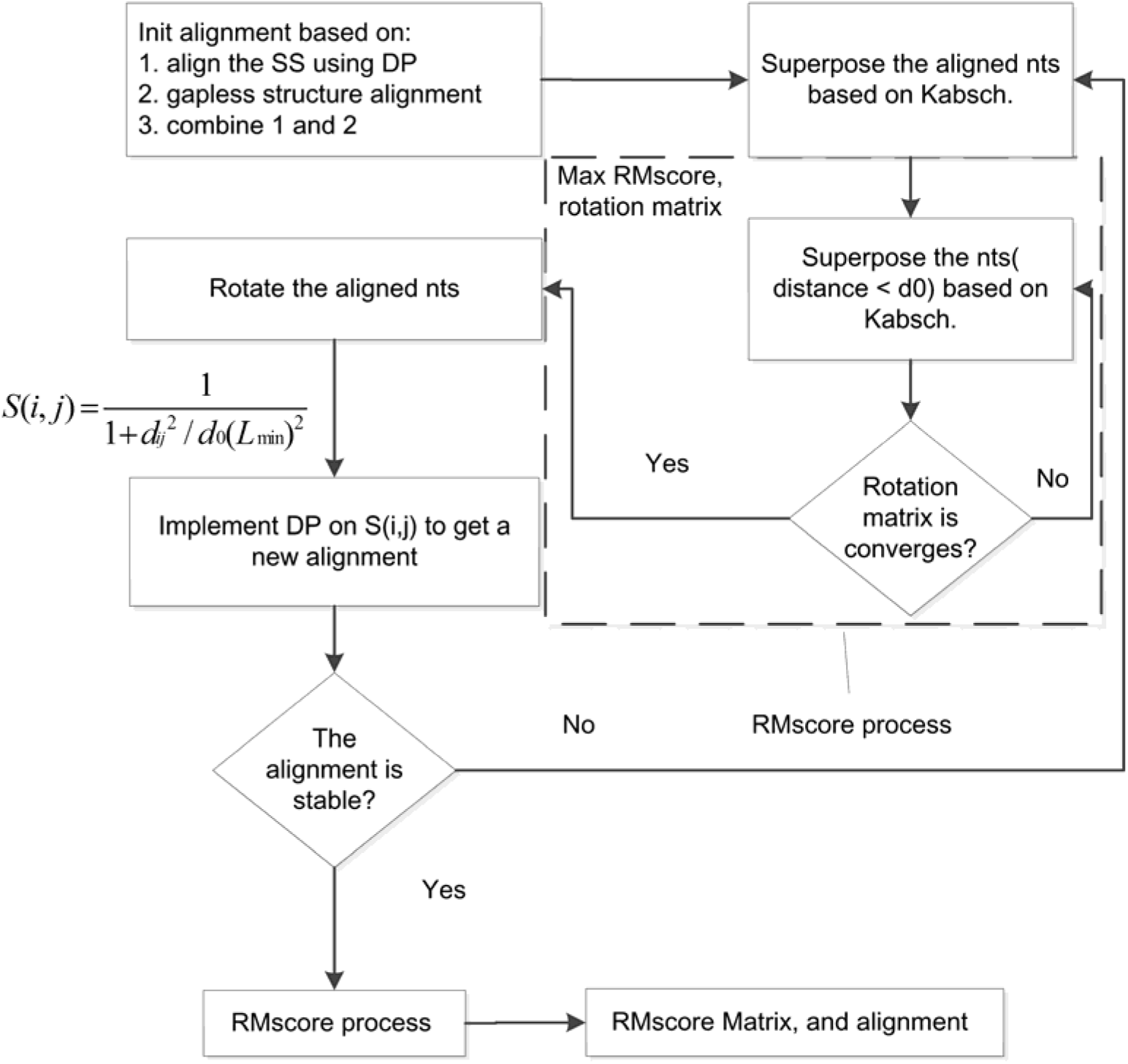
The flowchart of RMalign. SS stands for secondary structure and DP stands for dynamic programming algorithm. This process is a modification of TMalign.

**Figure S5.**
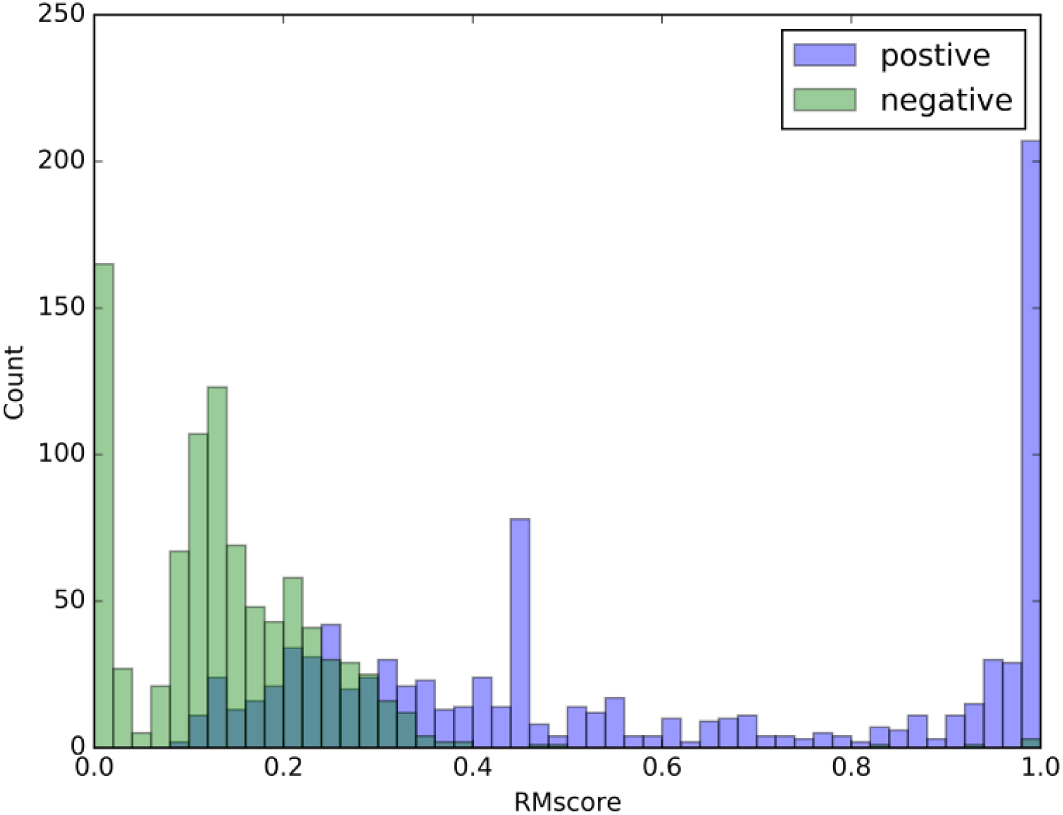
Distribution of RMscore benchmarking in balance-FSCOR. Positive pairs are RNA pairs with the same functions. Negative pairs are RNA pairs with different functions. The figure shows that most negative pairs have a lower RMscore and most positive pairs have a higher RMscore.

**Figure S6.**
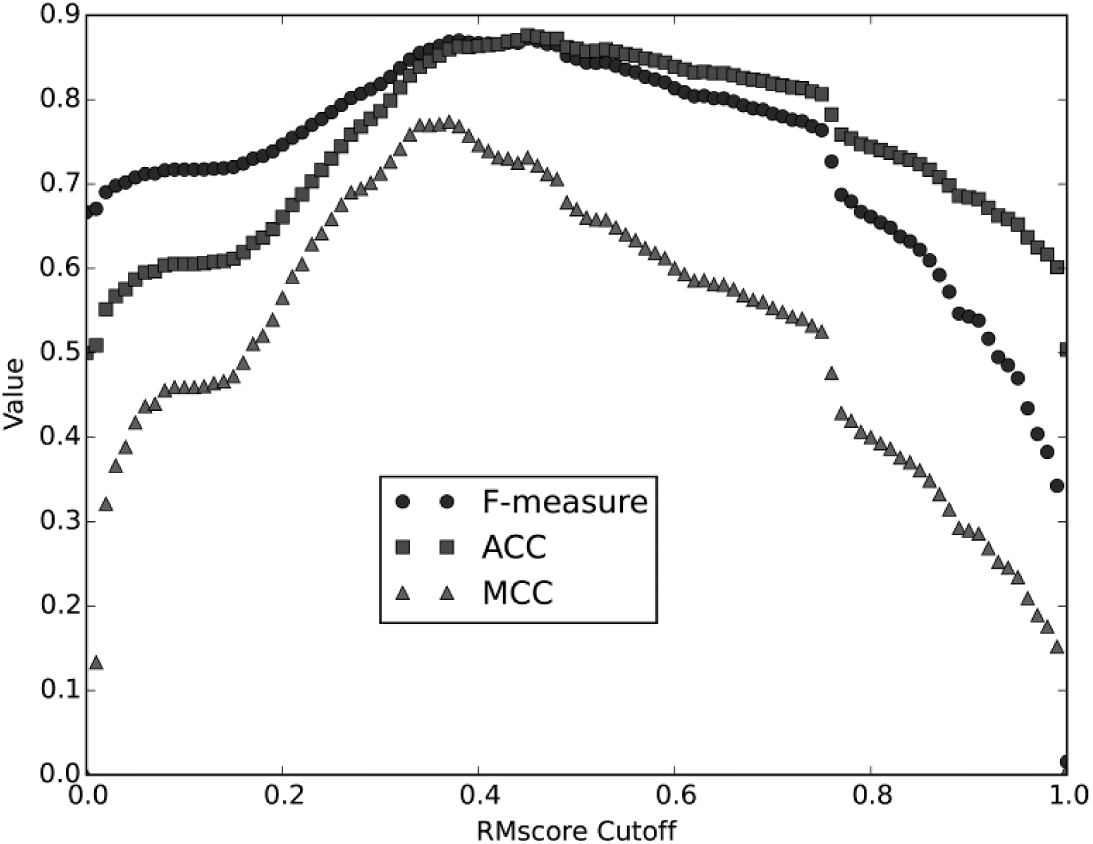
RMscore cut off vs F-measure, ACC, MCC. F-measure, ACC (accuary) and MCC are plotted against RMscore cut off selected to predict the positive or negative pairs when benchmarking on balance-FSCOR.

**Figure S7.**
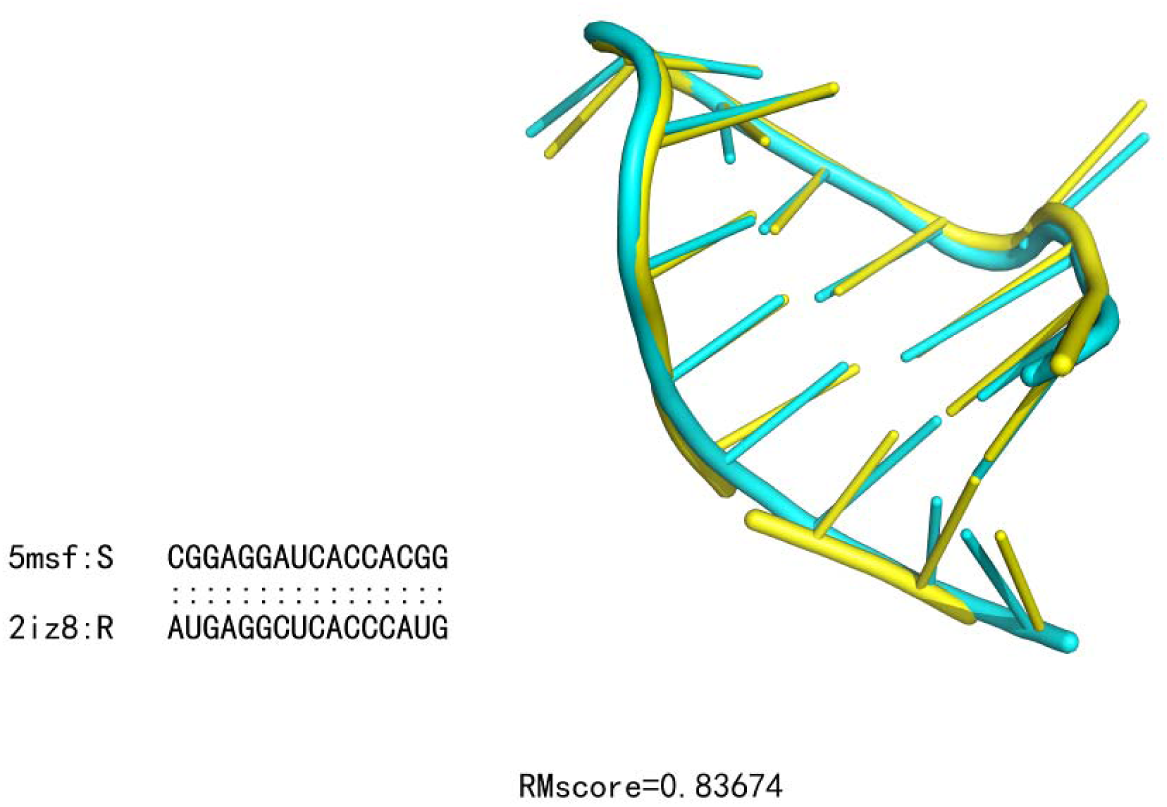
A false positive example of RMalign benchmarking in balance-SCOR. 5msf:S (yellow) is superposed on 2iz8:R (cyan) by RMalign. Two RNA structures are very similar (RMscore = 0.837), however, they are belonged to two distinct function classes in FSCOR (Phage_coat_protein_binding, MS2_phage_coat_protein_binding_stem-loop).

**Figure S8.**
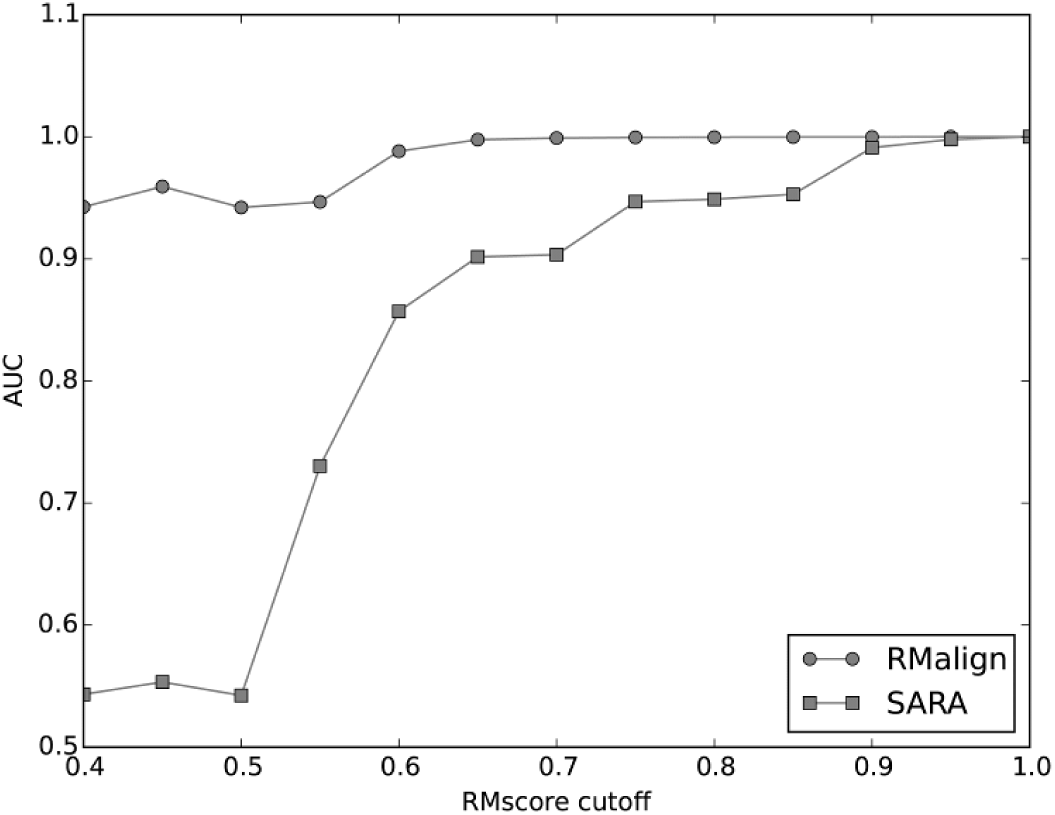
Benchmarking of RMalign and SARA on balance-x-FSCOR.

